# Combining multivariate genomic approaches to elucidate the comorbidity between ASD and ADHD

**DOI:** 10.1101/2020.04.22.054825

**Authors:** Hugo Peyre, Tabea Schoeler, Chaoyu Liu, Camille Michèle Williams, Nicolas Hoertel, Alexandra Havdahl, Jean-Baptiste Pingault

## Abstract

**Background:** Several lines of evidence point toward the presence of shared genetic factors underlying Autism Spectrum Disorder (ASD) and Attention Deficit Hyperactivity Disorder (ADHD). However, Genome-Wide Association Studies (GWAS) have yet to identify risk variants (i.e. Single-Nucleotide Polymorphisms, SNPs) shared by ADHD and ASD.

**Methods:** Two complementary multivariate analyses – genomic structural equation modelling (SEM) and colocalization analysis – were exploited to identify the shared SNPs for ASD and ADHD, using summary data from two independent GWAS of ASD (N=46,350) and ADHD individuals (N=55,374).

**Results:** Genomic SEM identified 7 novel SNPs shared between ASD and ADHD (*p*_genome-wide_<5e-8), including three SNPs that were not identified in any of the original univariate GWAS of ASD and ADHD (rs227378, rs2391769 and rs325506). We also mapped 4 novel genes (MANBA, DPYD, INSM1, and PAX1) to SNPs shared by ASD and ADHD, as well as 4 genes that had already been mapped to SNPs identified in either ASD or ADHD GWAS (SORCS3, XRN2, PTBP2 and NKX2-4). All the shared genes between ADHD and ASD were more prominently expressed in the brain than the genes mapped to SNPs specific to ASD or ADHD. Colocalization analyses revealed that 44% percent of the SNPs associated with ASD (*p*<1e-6) colocalized with ADHD SNPs and 26% of the SNPs associated with ADHD (*p*<1e-6) colocalized with ASD SNPs.

**Conclusions:** Using multivariate genomic analyses, the present study reveals the shared genetic pathways that underlie ASD and ADHD. Further investigation of these pathways may help identify new targets for treatment of these disorders.

## INTRODUCTION

Attention Deficit Hyperactivity Disorder (ADHD) and Autism Spectrum Disorder (ASD) are two common neurodevelopmental disorders (1) with a prevalence in children between 5 to 7% (2) and 1 to 2%, respectively (3). ADHD is characterized by symptoms of inattention, impulsivity, and hyperactivity, and ASD by a deficit in social communication as well as restricted and repetitive patterns of interests and behaviors. Although both disorders are distinctive in terms of diagnostic criteria, there is a considerable overlap in symptomatology. Individuals with ASD commonly display inattention, impulsivity, and hyperactivity symptoms (4) and likewise, individuals with ADHD often have impaired social and communication skills (5–7). In addition, ASD and ADHD are frequently comorbid; approximately one third of children with ASD are also diagnosed with ADHD (8,9) and one fifth of children with ADHD meet diagnostic criteria for ASD (7).

Little is known about what causes the association between ASD and ADHD. Both disorders are thought to be caused by a complex interplay between environmental and genetic risk factors (10,11). Shared environmental risk factors, such as preterm birth (12,13) or prenatal exposure to valproate (14,15) might partially explain the association between the disorders. Several lines of evidence also point toward the presence of shared genetic factors in ASD and ADHD. In family-based studies, relatives of children with ASD are at a higher risk for ADHD than relatives of children without ASD (16,17) and in twin studies, researchers report strong genetic correlations between traits related to ADHD and traits related to ASD (18,19). Moreover, a similar burden of rare protein-truncating variants has been found in individuals with ASD and those with ADHD (20). Most of the rare Copy Number Variants (CNVs) that are linked to ASD are also associated with ADHD (21,22). But less than 10% of ASD and ADHD liabilities can be accounted for by rare genetic variants (22,23). To this day, Genome-Wide Association Studies (GWAS) have not yet identified shared common risk variants (i.e. Single-Nucleotide Polymorphisms, SNPs) for ADHD and ASD (10,24,25). This is because the shared genetic risk has not yet been modelled using appropriate multivariate genomic approaches, such as genomic structural equation modelling (Genomic SEM) (26) or colocalization (27). Genomic SEM appears particularly useful to identify novel shared risk SNPs that remained undetected in univariate GWAS of overlapping traits (26,28), while colocalization allows identification of shared and specific SNPs after taking linkage disequilibrium (LD) into account. This latter method already elucidated the shared genetic risk between autoimmune diseases (29), lipid levels and cardiovascular outcomes (30), as well as schizophrenia and gene expression within human brain tissue (31,32).

As an alternative possibility, the association between ASD and ADHD might be explained by direct causal relationships between the two phenotypes, unidirectionally or bidirectionally. ADHD might lead to secondary impairments in social interaction and behavioral flexibility (33), while ASD might contribute to secondary attention problems, hyperactive and impulsive behaviours. These directional relationships could bias our interpretation of the colocalization analysis considering that a given SNP may colocalize between ADHD and ASD not because of shared genetic risk but because of the increase of ADHD symptoms in individuals with ASD or the increase of ASD symptoms in individuals with ADHD. To remedy to issue of directionality, the present study will be the first to conduct a Bidirectional Mendelian randomization (MR) analysis on ASD and ADHD GWAS to determine the effect of ADHD on ASD and the effect of ASD on ADHD (34,35).

This study used the aforementioned multivariate genomic approaches (Genomic SEM and colocalization) to identify SNPs shared by ASD and ADHD and SNPs specific to each disorder. Functional analyses were performed on shared and specific common genetic variants. Finally, a bidirectional Mendelian randomization analysis was performed to explore whether the shared genetic risk between ASD and ADHD was interpretable in terms of (reciprocal) causal relationships.

## METHODS AND MATERIALS

### GWAS Summary Statistics

Summary statistics for ASD and ADHD were obtained from the European ancestry subgroup of the Psychiatric Genomics Consortium and iPSYCH (PGC + iPSYCH). We used the most recent GWAS data on ASD (18,381 diagnosed ASD cases and 27,969 controls) (24) and ADHD individuals (20,183 diagnosed ADHD cases and 35,191 controls) (25).

### Genomic SEM

Genomic SEM (GenomicSEM R package) models the genetic covariance structure of GWAS summary statistics using LD score regression (LDSC) (26,36) to estimate the association of each SNP with the general factor (a latent variable corresponding to the shared variance between the ASD and ADHD GWAS). In the Genomic SEM, factor loadings were fixed to be equal between ASD and ADHD GWAS.

### Colocalization Analyses

Colocalization analyses were conducted on each genome-wide significant (*p*<5e-8) LD-independent (r2>0.2; window of 500kb) SNP associated with either ASD, ADHD, or the general factor to account for two types of false results when solely analysing individual SNP association parameters (*Supp. Figure 1*): (*i*), a SNP associated with a trait A may be falsely associated with the general factor of Trait A and B because a causal SNP of Trait B is in LD with the SNP of Trait A; (*ii*), the general factor might miss some SNPs that are in fact associated with both traits. For example, a shared causal SNP strongly associated with Trait A but less strongly associated with Trait B may erroneously not be linked to the general factor.

We used the R packages COLOC (37) and Hyprcoloc (38). Both methods implement a Bayesian test for the colocalization of two associations signals in a selected region (250kb around a SNP under study), using the summary statistics of two traits, to calculate five hypotheses: H0 (no association between the SNP and either trait), H1 (SNP association with trait 1 only), H2 (SNP association with trait 2 only), H3 (SNP association with both traits but not colocalizing, i.e. two distinct SNPs in LD); and H4 (SNP association with both traits and colocalizing). We set the prior probability that a SNP is causal in each trait to be identical (1e-4 is the recommended threshold for colocalization analysis in the context of GWAS (27,37,39)) and the prior probability that a SNP is causal for both traits at 5e-6 (meaning that 5 out of 100 SNPs that are associated with one trait are also associated with the other). A given SNP was considered to colocalize if H4>0.5 using COLOC or Hyprcoloc packages (31,40). To estimate the percentage of SNPs that colocalized between both traits more precisely, we performed a similar analysis but at the *p*-value threshold of 1e-6.

SNPs colocalizing between ADHD and ASD and associated with the general factor at p<5e-8 were considered as SNPs shared by ASD and ADHD. Among the SNPs not colocalizing between ADHD and ASD, those associated with a *p*-value below 5e-8 with ASD were classified as SNPs specific to ASD and those associated with a *p*-value below 5e-8 with ADHD were classified as SNPs specific to ADHD.

### Functional Analysis

Functional annotation and analyses were performed separately on SNPs shared by ASD and ADHD (i.e. SNPs colocalizing between ADHD and ASD), and SNPs specific to each disorder using FUMA (Functional Mapping and Annotation of Genome-Wide Association Studies) (http://fuma.ctglab.nl/) (41). Using FUMA SNP2GENE, SNPs are mapped to genes based on physical position (positional mapping of deleterious coding SNPs (CADD score≥12.37) and eQTL associations (we used 13 brain tissue types from the GTEx.v8 project). Moreover, SNPs are mapped to genes based on 3D chromatin interactions (chromatin interaction mapping; Hi-C data of 2 tissue types: adult and foetal cortex (42)). Chromatin interaction mapping can involve long-range interactions. GENE 2FUNC then annotates these genes in a biological context. Among the other functions of GENE 2FUNC, tissue specific expression patterns (based on GTEx v6 RNA-seq data) for each gene were visualized as an interactive heatmap and globally for SNPs that are shared by ASD and ADHD and for those that are specific to ASD and ADHD.

### Bidirectional Mendelian Randomization Analyses

We ran bidirectional summary data Mendelian randomization (MR) analysis to determine the effect of the liability for ADHD on ASD and the liability for ASD on ADHD (34,35,43). LD-independent SNPs (r2>0.001; window of 500kb) associated with ASD (n_SNPs_=15, with p<1e-6) were selected as instruments for ASD to estimate the effect of ASD on ADHD. Associations were also ascertained in the opposite direction, using n=41 SNPs (*p*<1e-6) as instruments for ADHD. We estimated the main effects using the inverse weighted variance (IVW) estimator, which consists of a linear regression of the instrument-outcome association estimates on the instrument-exposure association estimates, weighted by the inverse of the variance of the instrument-outcome association estimates. A number of MR methods were also employed for sensitivity analyses to examine potential violations of the MR assumptions, such as exclusion restriction assumption due to pleiotropy (i.e. the SNP used as instrument does not affect directly both exposure and outcome), including the weighted median-based and mode-based because they allow some genetic instruments (i.e. SNPs) to be invalid (44). We also conducted leave-one-out sensitivity analyses to check for a disproportionate influence of individual SNPs on overall effect estimates using the IVW method. We additionally applied MR-Egger regression (45,46) to estimate the intercept (corresponding to a test of directional pleiotropy) in addition to the slope in the regression. The I²_GX_ statistic was used to test for heterogeneity in MR Egger regression.

Because only few SNPs associated with ASD and ADHD at genome-wide significance level (*p*<5e-8) are available in the current GWAS (Grove et al. (24), Demontis et al. (25)), MR analyses were conducted at a *p*-value threshold of 1e-6. As MR analysis were performed with instruments below the conventional GWAS threshold (i.e. *p*<5e-8) we also applied the Mendelian randomization robust adjusted profile score (MR-RAPS) to account for weak instrument bias (47).

## RESULTS

### Genomic SEM

Genomic SEM was conducted on 6,971,687 SNPs that were present in both ASD (number of available SNPs=7,757,027) and ADHD (number of available SNPs=8,094,094) GWAS. There were 232 genome-wide significant (*p*<5e-8) SNPs identified in the general factor GWAS, including 7 LD-independent SNPs following clumping. Two of the 7 SNPs (rs1222063 and rs4916723) were previously reported in the ADHD GWAS from which we obtained the summary statistics (**Table 1**). The Manhattan plot of the general factor GWAS is depicted in **Figure 1**.

**Table 1.**
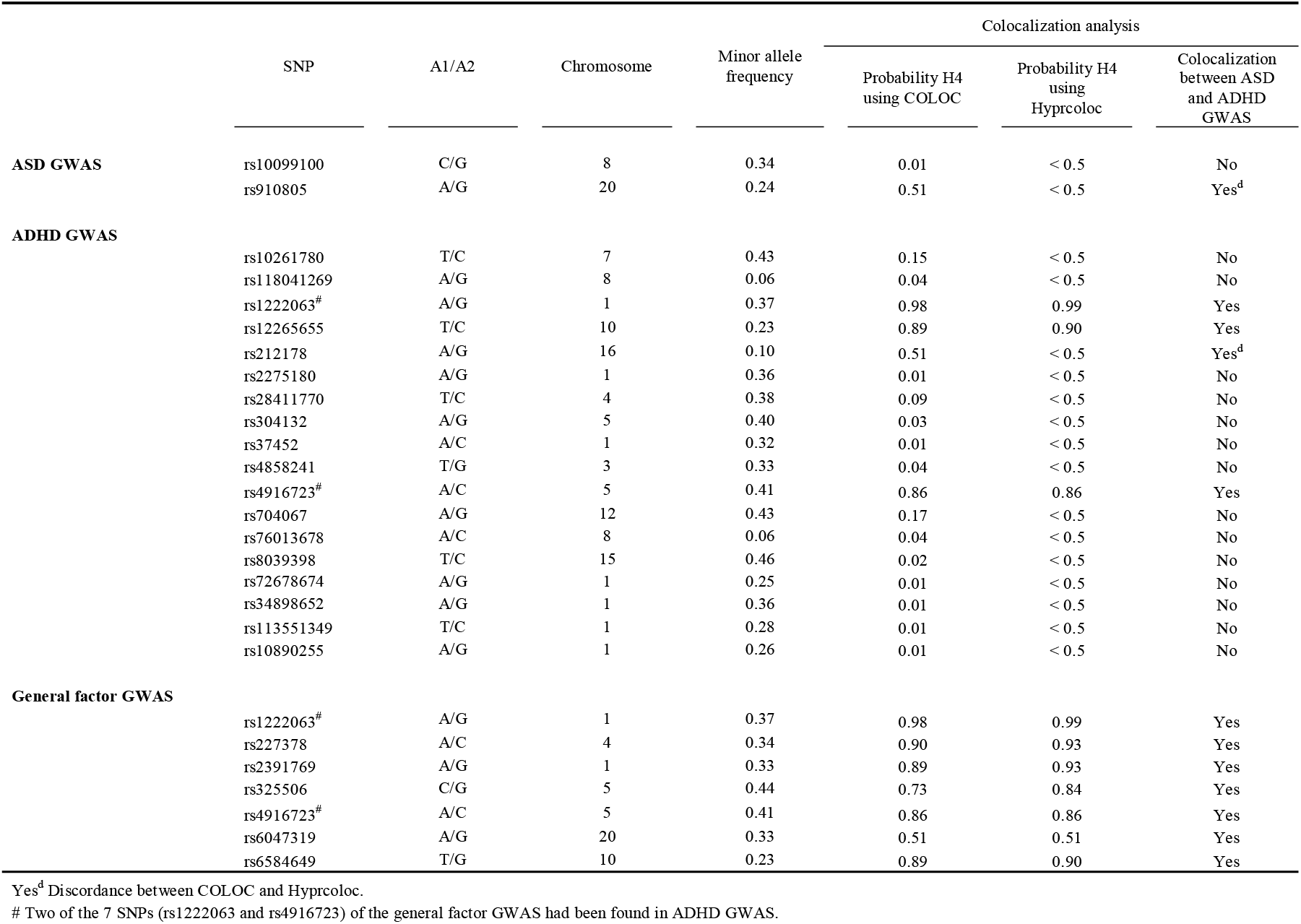
Colocalization analysis of SNPs of ASD, ADHD and the general factor (using COLOC and Hyprcoloc).

**Figure 1.**
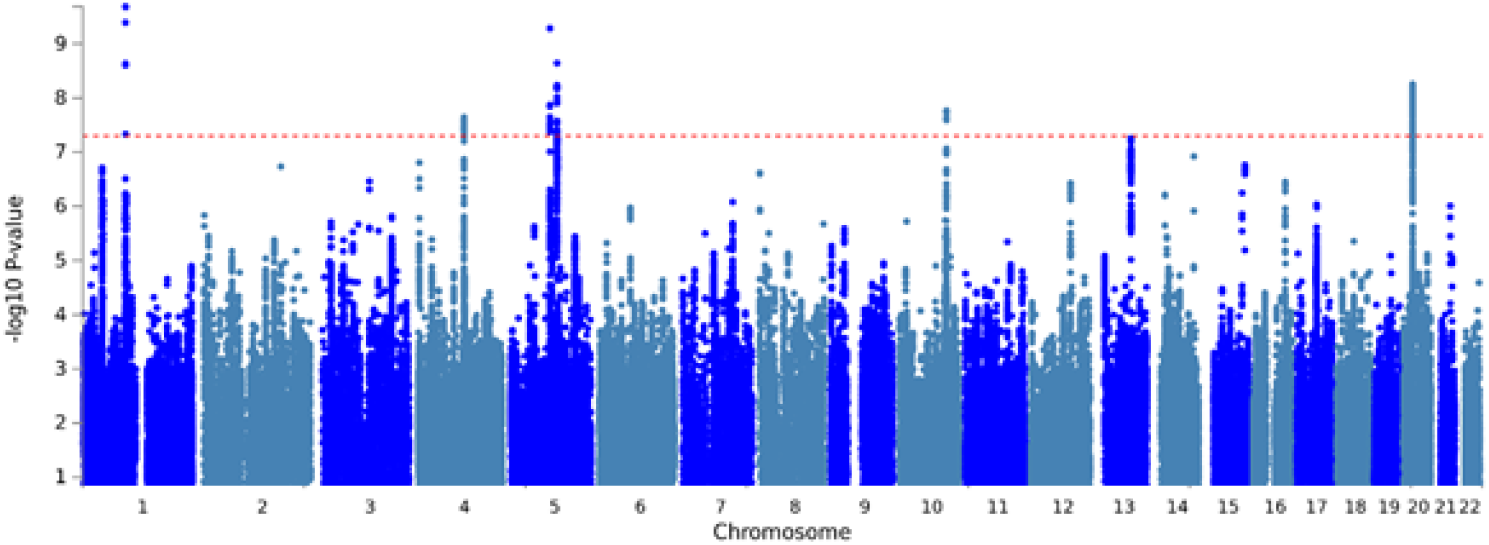
Manhattan plot of the general factor of ASD and ADHD using Genomic SEM.

### Colocalization Analyses

The main results of the colocalization analysis are summarized in **Table 1**. One out of the two SNPs associated with ASD colocalized with ADHD, 4 out of 18 SNPs associated with ASD colocalized with ADHD, and all the 7 SNPs associated with the general factor colocalized between ASD and ADHD GWAS.

After clumping, we found two SNPs associated with ASD (and 16 when *p*-value threshold was set at 1e-6) and 18 SNPs associated with ADHD (and 50 when *p*-value threshold was set at 1e-6). Among the two SNPs (rs10099100 (*Supp. Figure 2*) and rs910805 (*Supp. Figure 3*)) associated with ASD after clumping, one (rs910805) colocalized with ADHD GWAS. When *p*-value was set at 1e-6, the proportion of SNPs of ASD that colocalized with ADHD GWAS was 44% (7/16). Among the 18 SNPs associated with ADHD, 4 of them colocalized with ASD GWAS. When *p*-value threshold was set at 1e-6, the proportion of SNPs of ASD that colocalized between ADHD GWAS was 26% (13/50). Among the 7 SNPs associated with the general factor, all of them colocalized between ADHD and ASD GWAS (**Figure 2** and *Supp. Figures 4-9*). Three of them did not reach statistical significance in either the ASD and ADHD GWAS: rs227378, rs2391769, and rs325506 and may be considered as new SNPs for the shared genetic liability between ASD and ADHD. When the *p*-value threshold was set at 1e-6, the proportion of SNPs shared by ASD and ADHD as determined by Genomic SEM that colocalized between ASD and ADHD GWAS was 79.2% (19/24).

**Figure 2.**
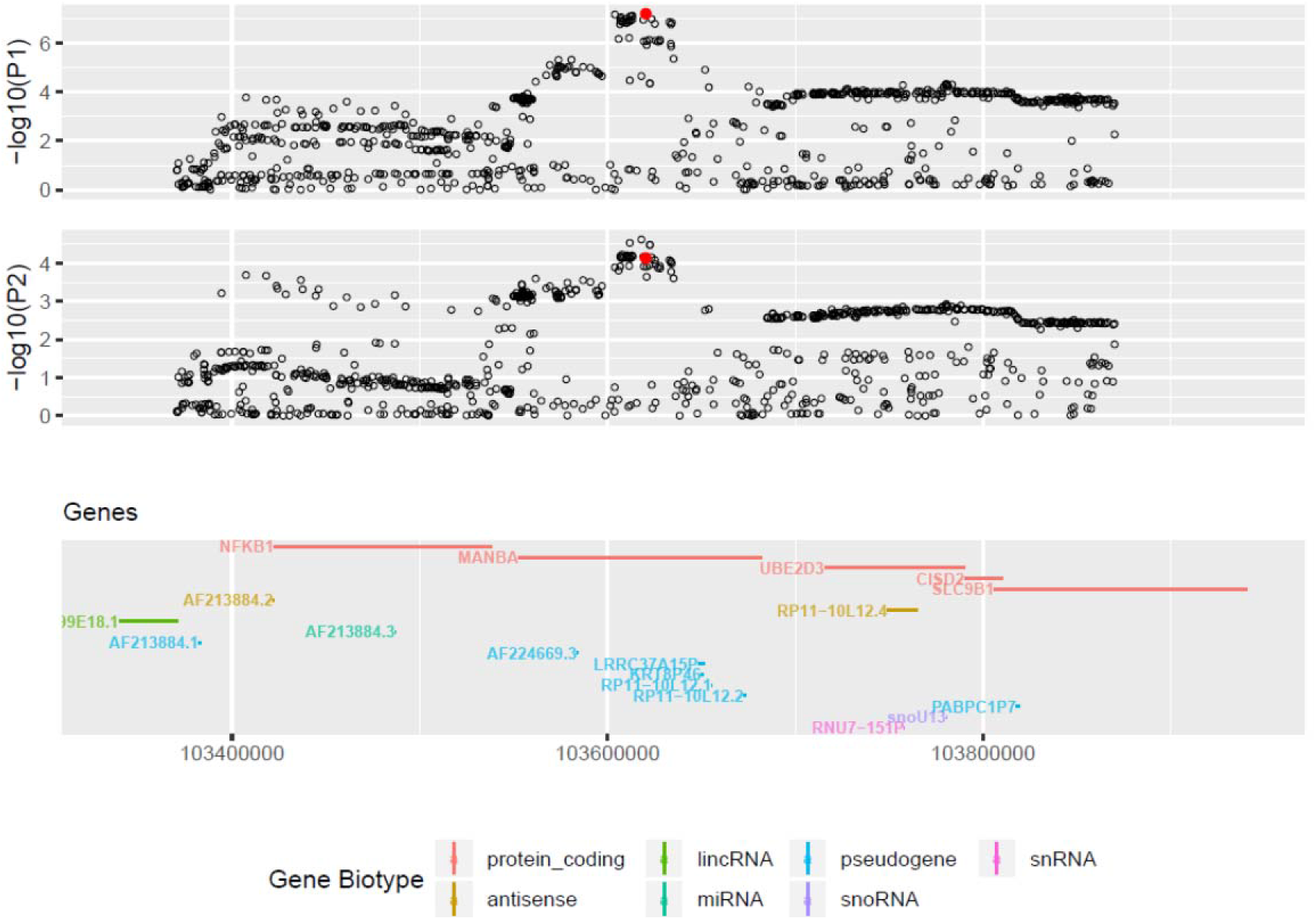
Region around rs227378 (250kb around the SNP) in ADHD (upper figure [P1]) and ASD (lower figure [P2]) GWAS.

### Functional Analyses of SNPs shared by ASD and ADHD

We applied FUMA to the 7 SNPs of the general factor. These 7 SNPs mapped onto 8 genes, with 4 that correspond to novel genes of ASD and ADHD: MANBA, DPYD, INSM1, and PAX1.

Positional and eQTL mapping of deleterious coding SNPs prioritized three genes: XRN2 in region 20p11.22 around rs6047319 for XRN2 (*Supp. Figure 8*), SORCS3 in region 10q25.1 around rs6584649 (*Supp. Figure 9*), and MANBA in region 4q24 around rs227378 (**Figure 2**). The prioritized three genes included two genes which were previously reported as candidates in the study conducted on ASD by Grove et al. (24) (XRN2) and the one conducted on ADHD by Demontis et al. (25) (SORCS3), while MANBA was a novel gene. Gene Ontology annotations related to SORCS3 include neuropeptide receptor activity (SORCS3 showed a specific expression in brain; *Supp. Figure 10 (A)*). Mutations in the gene MANBA are associated with beta-mannosidosis, a lysosomal storage disease that has a wide spectrum of neurological implications (48). Five Supplementary genes were identified by chromatin interaction mapping: PTBP2 and DPYD (in region 1p21.3 around rs2391769), INSM1 (in region 20p11.23 around rs6047319), and NKX2-4 and PAX1 (in region 20p11.22 around rs6047319) (*Supp. Table 1*). Among them, PTBP2 and NKX2-4 were previously reported as candidates in the study conducted on ASD by Grove et al. (24) (and (49)), while DPYD, INSM1, and PAX1 were novel genes. 1p21.3 microdeletions affecting DPYD have been identified in individuals with ASD (50), intellectual disability (51) and has been considered as a risk loci for schizophrenia (52,53). INSMI is known to play a key role in neurogenesis and neuroendocrine cell differentiation during embryonic and/or fetal development and NKX2-4 showed a specific expression in hypothalamus (*Supp. Figure 10 (A)*). Some SNPs shared by ASD and ADHD did not map to a gene (rs1222063, rs325506, rs4916723). However, GWAS data (*Supp. Data 1* for more references) suggest that the region around rs325506 (5q21.2) was associated with several traits including depression (54) and educational achievement (55), and that the region around rs4916723 (5q14.3) was associated with neuroticism and alcohol-consumption (56,57). Globally SNPs shared by ASD and ADHD were mapped to genes with specific expression in the brain (**Figure 3**).

**Figure 3.**
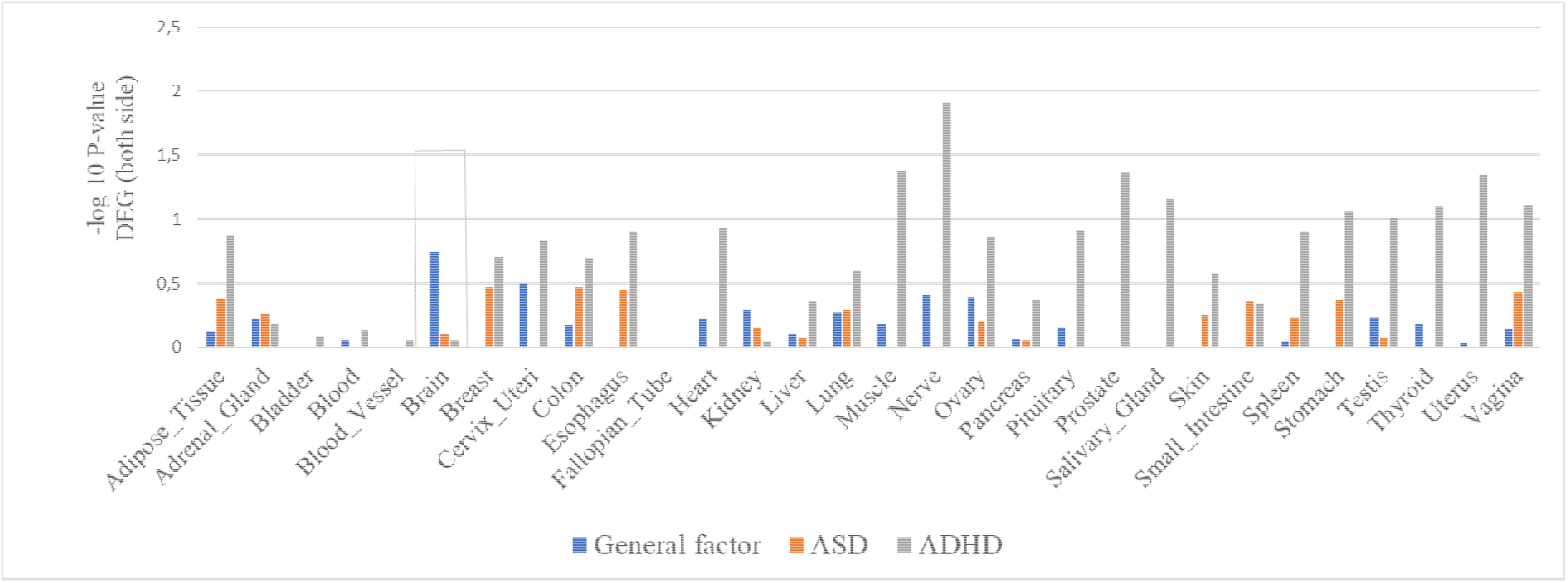
Tissue specific expression patterns (average of normalized expression per label) using GTEx v6 RNA-seq data (FUMA GENE2FUNC) of the SNPs shared by ASD and ADHD and those that are specific to ASD and ADHD.

### Functional Analyses of SNPs Specific to ASD

We applied FUMA to the SNP specific to ASD (rs10099100). Positional mapping of deleterious coding SNPs prioritized two genes (SOX7 and RP1L1) which were previously reported as candidates by Grove et al. (24) and Alonso-Gonzalez et al. (49). These genes showed a low brain-specific expression (**Figure 3** and *Supp. Figure 10 (B)*).

### Functional Analyses of SNPs Specific to ADHD

We applied FUMA to the 14 genome-wide significant SNPs specific to ADHD. Positional mapping of deleterious coding SNPs (22 genes), eQTL mapping (5 genes), and chromatin interaction mapping (27 genes) prioritized 37 unique genes (some genes identified by multiple mapping methods; *Supp. Table 3*). We used different eQTL and chromatin mapping parameters on FUMA than the study by Demontis et al. (25). Therefore only 15 out of the 34 genes were previously reported as candidate genes by Demontis et al. (25). Each SNPs rs8039398, rs28411770, and rs704067 were mapped to a single prioritized gene (respectively SEMA6D, PCDH7 and DUSP6) and were therefore highly likely to drive the association signal. Tissue expression of the 37 genes did not indicate a clear brain-specific expression (**Figure 3**).

### Bidirectional Mendelian Randomization Analysis

Using bidirectional Mendelian randomization, we found that the risk of ASD was associated with an increased risk of ADHD across all MR approaches (β – MR-IVW=0.51 (0.09), *p*-value<0.001, number of SNPs=15; **Figure 4** and *Supp. Table 4*). Testing the reverse direction, we found that the risk of ADHD was associated with an increased risk of ASD (β – MR-IVW=0.34 (0.05), *p*-value<0.001, number of SNPs=41) across most MR approaches. MR-Egger implicated an effect from ASD to ADHD (β=0.76 (0.34), *p*-value=0.025, I²_GX_ statistic=96.0%) but no significant effect in the reverse direction (ADHD to ASD, β=0.06 (0.21), *p*-value=0.8, I²_GX_ statistic=96.1%).

**Figure 4.**
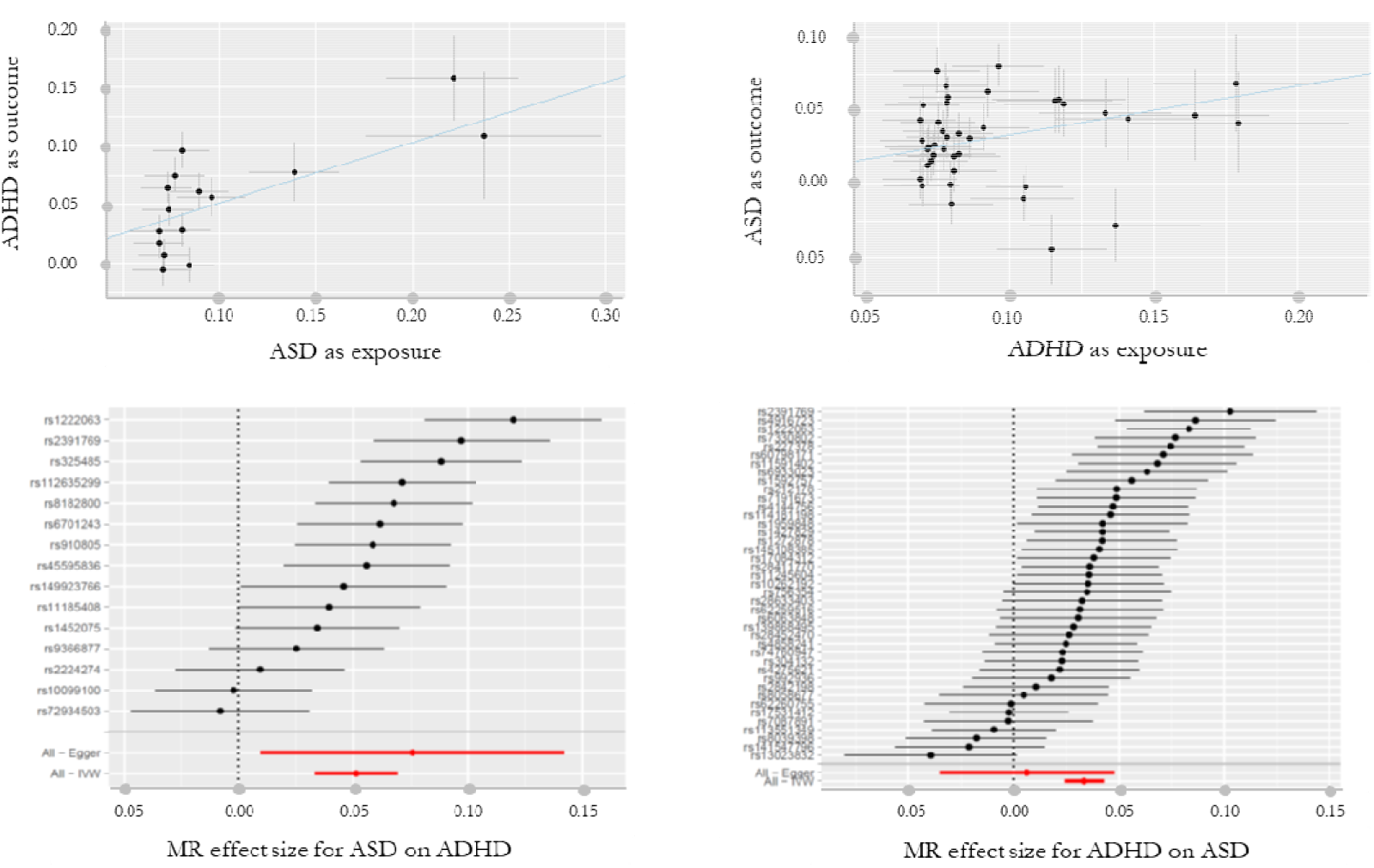
Bidirectional MR using the inverse weighted variance (IVW) estimator. Causal relationship between ASD as exposure (*p*-value at 1e-6) and ADHD as outcome (left). Causal relationship between ADHD as exposure (*p*-value at 1e-6) and ASD as outcome (right).

Leave-one-out sensitivity analyses did not indicate a disproportionate influence of an individual SNP in any of our MR-IVW analyses. MR-Egger did not indicate directional pleiotropy in both directions.

## DISCUSSION

Using powerful multivariate genomic approaches, we examined the joint genetic architecture of ASD and ADHD. To date, none of the SNPs identified as genome-wide significant in the univariate GWAS of ASD and ADHD were shared between the two disorders. Implementing genomic structural equation modelling, we identified 7 novel shared risk SNPs between ASD and ADHD. Among them, three were not identified in either of the original univariate GWAS of ASD and ADHD (rs227378, rs2391769 and rs325506). We also mapped 4 novel genes (MANBA, DPYD, INSM1, and PAX1) to SNPs shared by ASD and ADHD, as well as 4 genes that had been already mapped to SNPs identified in either ASD or ADHD GWAS (SORCS3, XRN2, PTBP2 and NKX2-4). Globally, these 8 genes revealed a specific expression in the brain (**Figure 3**). MANBA and DPYD were mapped to novel SNPs (rs227378 and rs2391769 respectively). Mutations in these two genes have previously associated to neurodevelopmental disorders (48,50,51). The SNP rs2391769 was mapped to DPYD but also to PTBP2 in the genomic loci 1p21. However, the SNP rs227378 was mapped to a single prioritized gene (MANBA) and was highly likely to drive the association signal in the genomic loci 4q24.

Findings from the present study critically add to previous univariate GWAS studies not only by identifying novel shared SNPs but also by identifying those suspected to be specific to one or the other disorder. The pattern of findings appeared strikingly different between shared and specific SNPs. Only one SNP (rs10099100) was considered specific to ASD. This SNP mapped two genes (SOX7 and RP1L1) with a low expression specificity for brain tissues (SOX7 is mostly expressed in lung and RP1L1 in the retina). Similarly, genes mapped to ADHD specific SNPs did not show a clear expression specific to brain tissues (**Figure 3**). Our analyses therefore suggest that identifying genetic variants with specific effects on either disorder may be less straightforward. The identification of specific genetic and environmental factors of ASD and ADHD remains crucial to uncover the complex aetiology of these neurodevelopmental disorders and should be the focus of further investigations (58).

Using colocalization analysis – a well-suited method to identify shared genetic risks between two traits (29–32) – we confirmed that all the 7 SNPs identified with genomic structural equation modelling were indeed shared between the two disorders. In addition, a substantial part of SNPs for ASD were associated with ADHD and vice versa. We found that 44% percent of the SNPs associated with ASD colocalized with ADHD SNPs, and 26% of the SNPs associated with ADHD colocalized with ASD SNPs (at *p*<1e-6). These results are consistent with the abundant evidence for shared genetic factors in ASD and ADHD from family-based (16,17), twin-based (18,19), and clinical studies on rare genetic variants (20– 22). In addition, results from bidirectional Mendelian randomisation analyses indicated that ASD increased the risk of ADHD and vice versa. These results suggest that ADHD leads to secondary impairments in social interaction and behavioral flexibility and that ASD contributes to secondary attention problems, hyperactive and impulsive behavior.

### Limitations

Although summary statistics for ASD and ADHD were obtained from the largest samples available, they enable the identification of only a few SNPs at the conventional genome-wide *p*-value threshold of 1e-8 (and thus few genes, especially for ASD). This contrasts with the findings that many more genes have been linked to ASD or ADHD with regards to rare genetic variants (20–22). GWAS of ASD and ADHD based on larger samples are needed to explore the genetic complexity of these disorders. Yet, the results of our colocalization were not affected by the number of SNPs. In fact, our estimates of the proportion of ASD associated SNPs that colocalize with ADHD associated SNPs and the proportion of ADHD associated SNPs that colocalize with ASD associated SNPs were very similar at the conventional *p*_genome-wide_<5e-8 (50% and 22% respectively) and at *p*<1e-6 (44% and 26%, respectively).

## Conclusion

Our findings support that ADHD and ASD, two of the most common neurodevelopmental disorders, share several common genetic risk variants. Our study revealed that about about half of the common genetic variants associated with ASD are also linked to ADHD, and about one quarter of the common genetic variants associated with ADHD are also linked to ASD. Using powerful multivariate genomic approaches, we identified 7 novel SNPs shared between ADHD and ASD, including three SNPs not identified in the univariate GWAS of ASD and ADHD, and mapped 4 novel genes (MANBA, DPYD, INSM1, and PAX1) to SNPs shared by ASD and ADHD. Further investigation of these pathways may help identify new targets for treatment of these disorders.

## Supporting information

Dupp Data

## Acknowledgement

JBP is supported by the Medical Research Foundation Emerging Leaders 1st Prize 2018 Adolescent Mental Health.

