## Supplementary material for "Combining multivariate genomic approaches to elucidate the comorbidity between ASD and ADHD": Supp Tables and Figures

**Supp. Figure 1. Conceptual framework of colocalization analysis.** Approaches taking solely individual SNP association parameters into account that did not consider Linkage Disequilibrium (LD) and may lead to false results in two ways (panels 1 and 2).

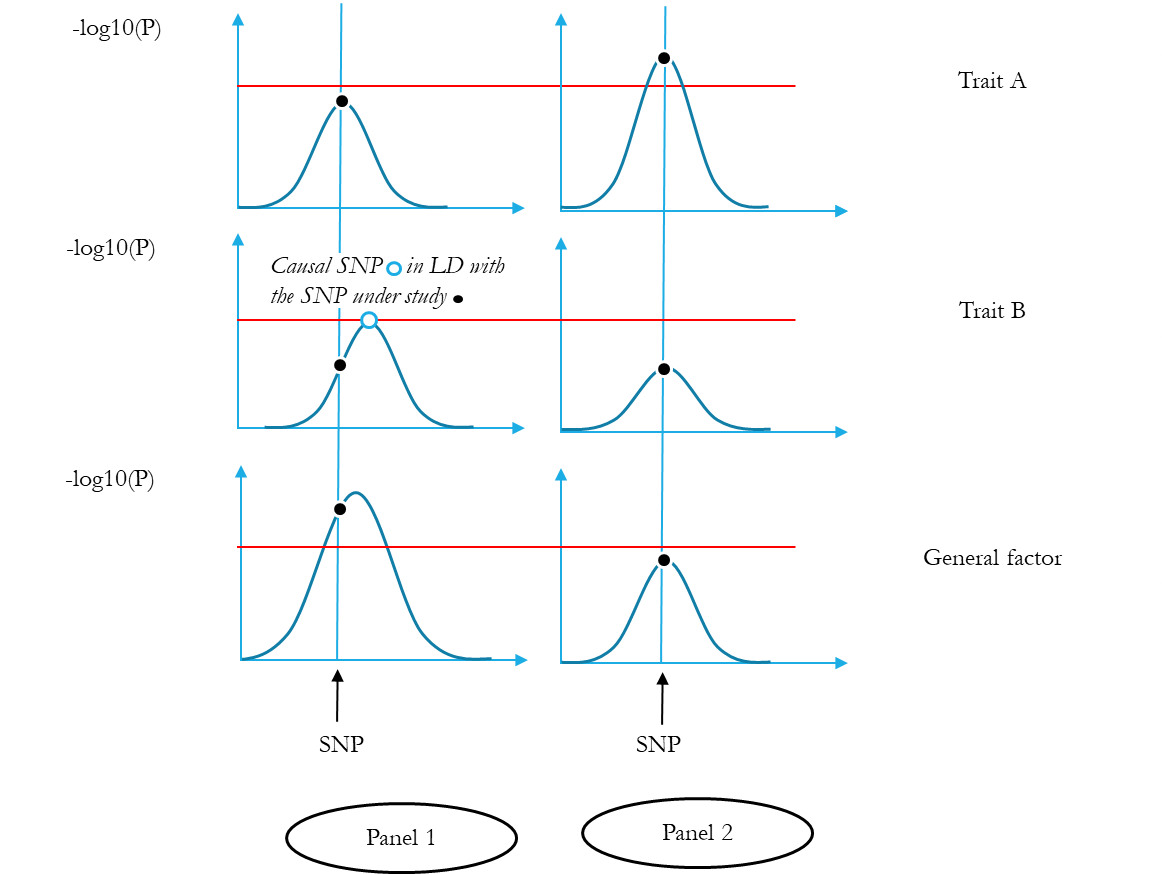

**Supp. Figure 2.** Region around rs10099100 (250kb around the SNP) in ADHD (upper figure [P1]) and ASD (lower figure [P2]) GWAS.

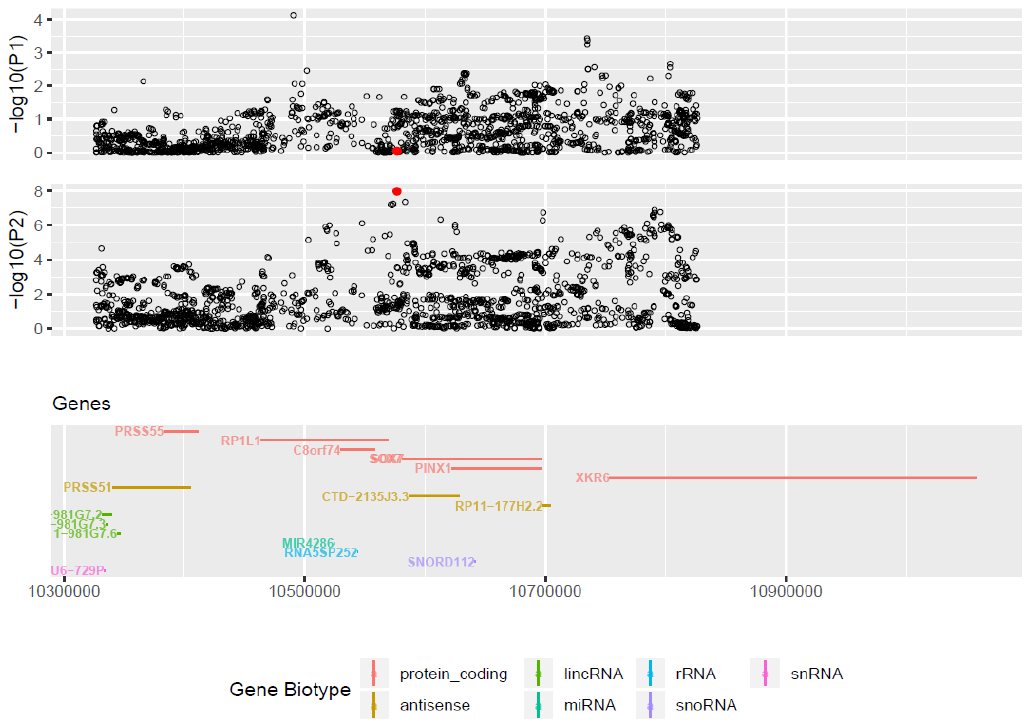

**Supp. Figure 3.** Region around rs910805 (250kb around the SNP loci) in ADHD (upper figure [P1]) and ASD (lower figure [P2]) GWAS.

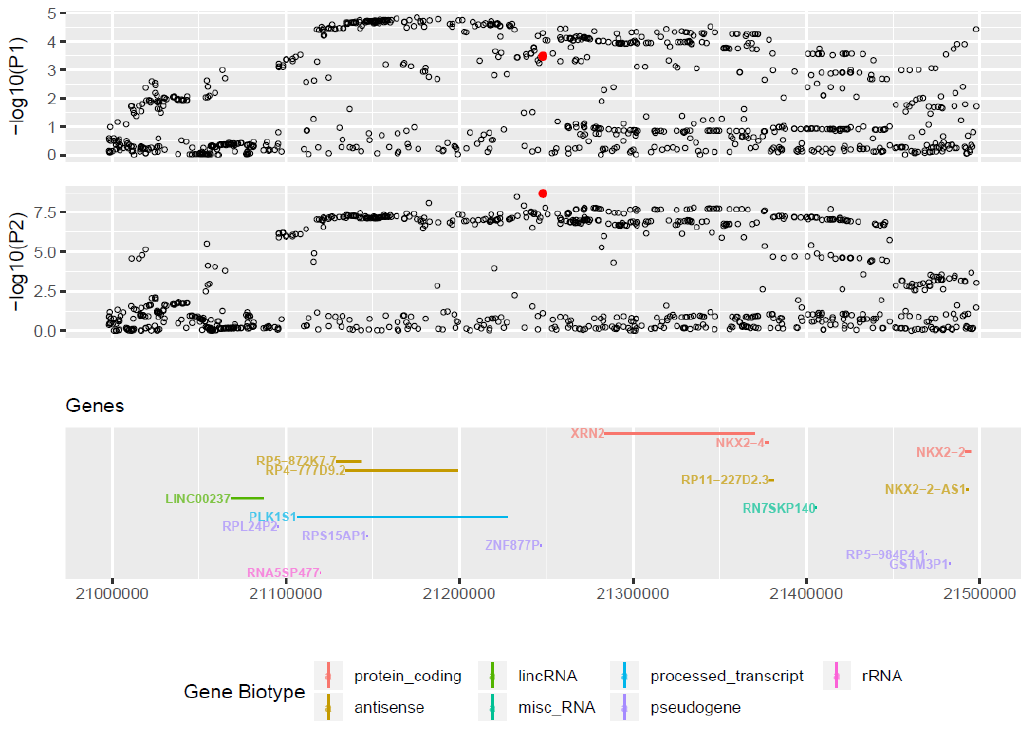

**Supp. Figure 4.** Region around rs1222063 (250kb around the SNP) in ADHD (upper figure [P1]) and ASD (lower figure [P2]) GWAS.

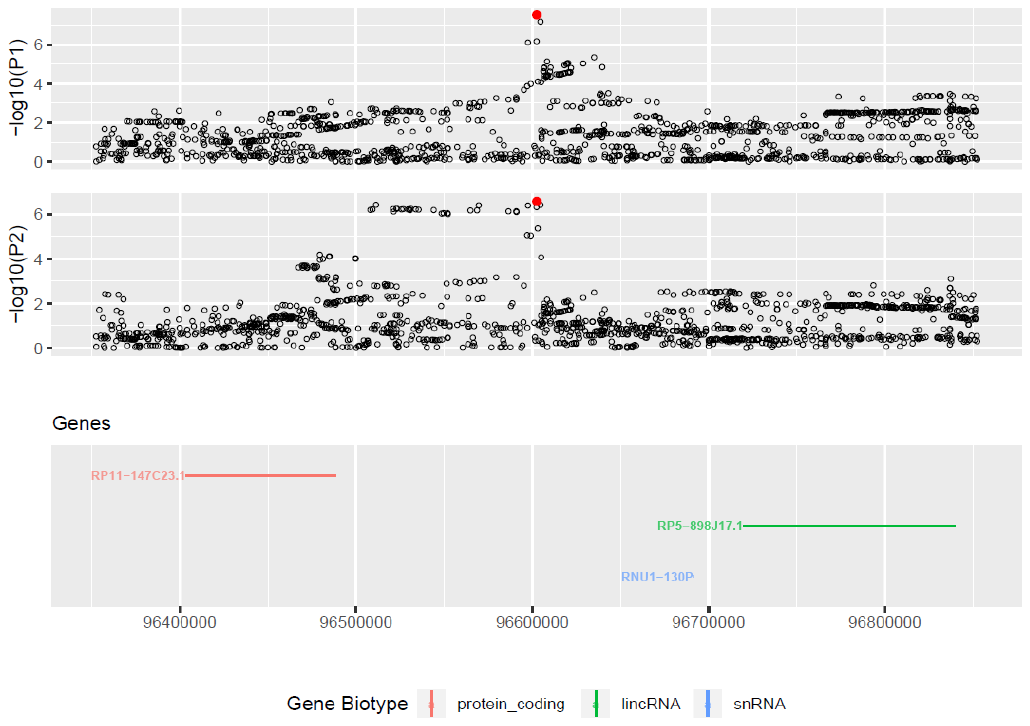

**Supp. Figure 5.** Region around rs2391769 (250kb around the SNP) in ADHD (upper figure [P1]) and ASD (lower figure [P2]) GWAS.

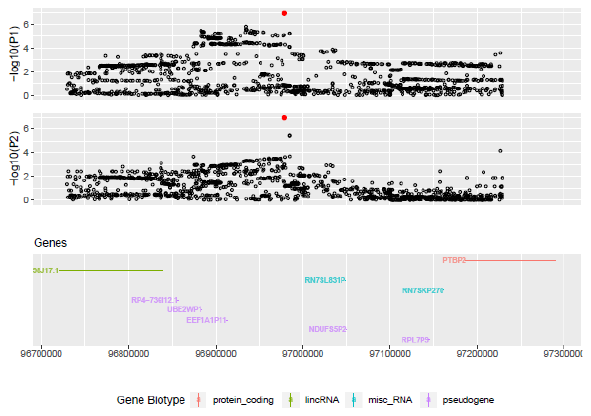

**Supp. Figure 6.** Region around rs325506 (250kb around the SNP) in ADHD (upper figure [P1]) and ASD (lower figure [P2]) GWAS.

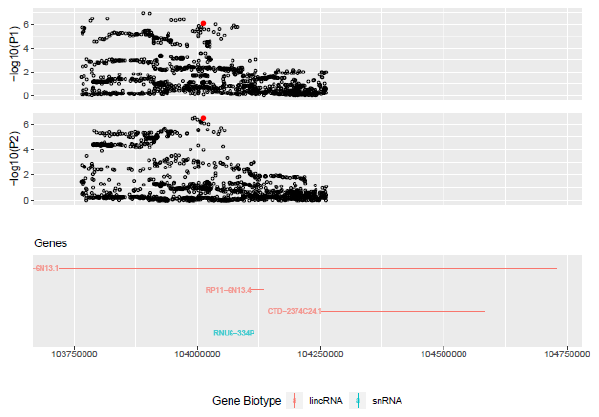

**Supp. Figure 7.** Region around rs4916723 (250kb around the SNP) in ADHD (upper figure [P1]) and ASD (lower figure [P2]) GWAS.

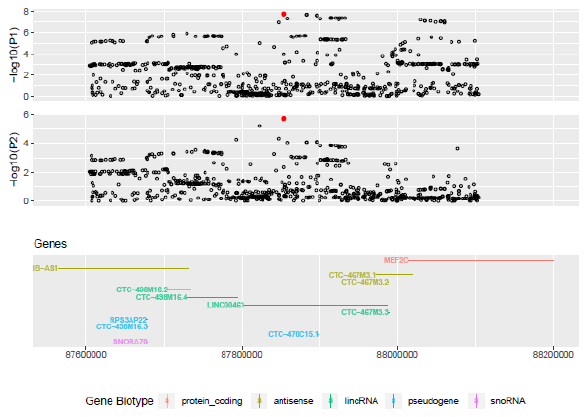

**Supp. Figure 8.** Region around rs6047319 (250kb around the SNP) in ADHD (upper figure [P1]) and ASD (lower figure [P2]) GWAS.

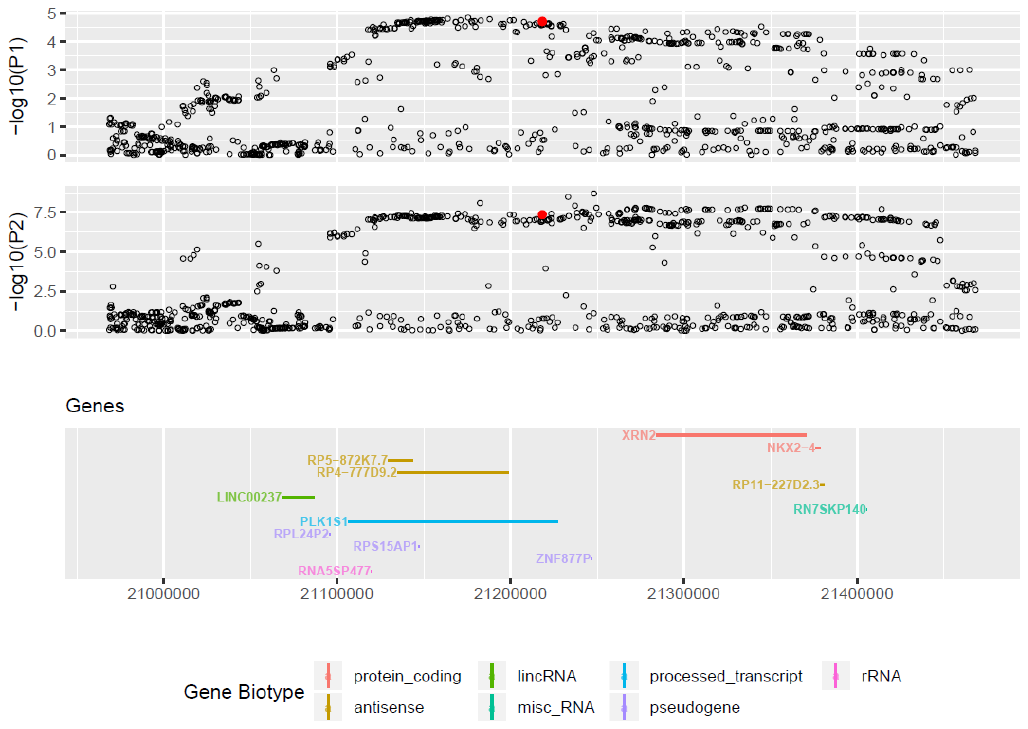

**Supp. Figure 9.** Region around rs6584649 (250kb around the SNP) in ADHD (upper figure [P1]) and ASD (lower figure [P2]) GWAS.

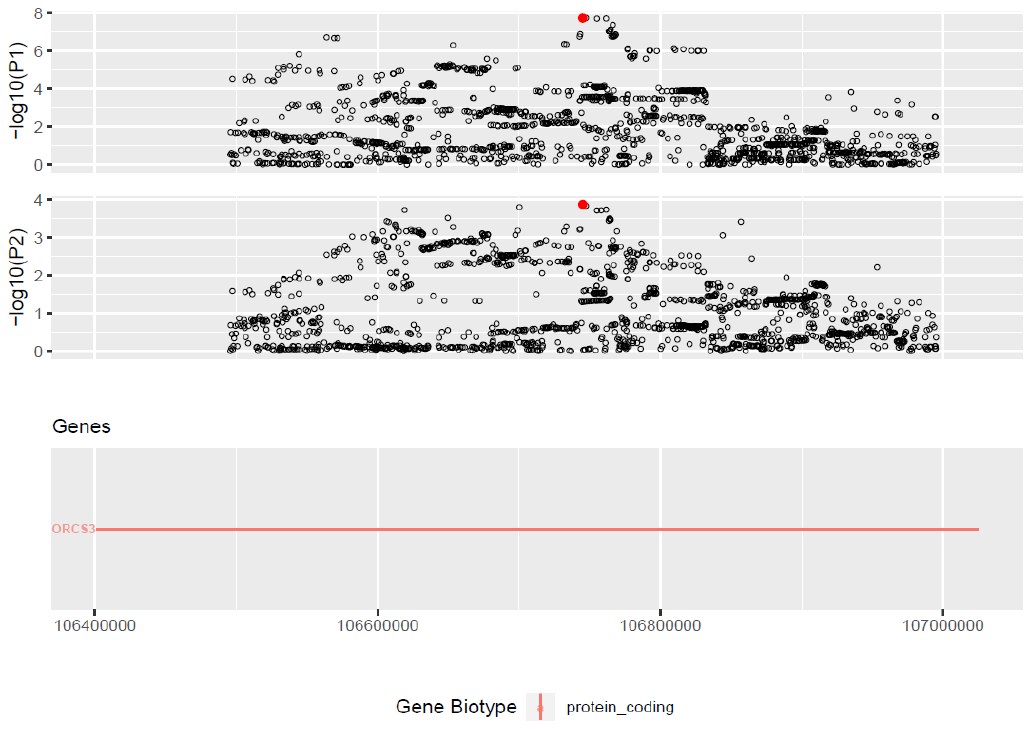

**SORCS3**

**Supp. Figure 10.** Tissue specific expression patterns (average of normalized expression per label) using GTEx v6 RNA-seq data (FUMA GENE2FUNC - gene expression heatmap) of SNPs that are **(A)** shared by ASD and ADHD and those that are specific to **(B)** ASD and **(C)** ADHD.

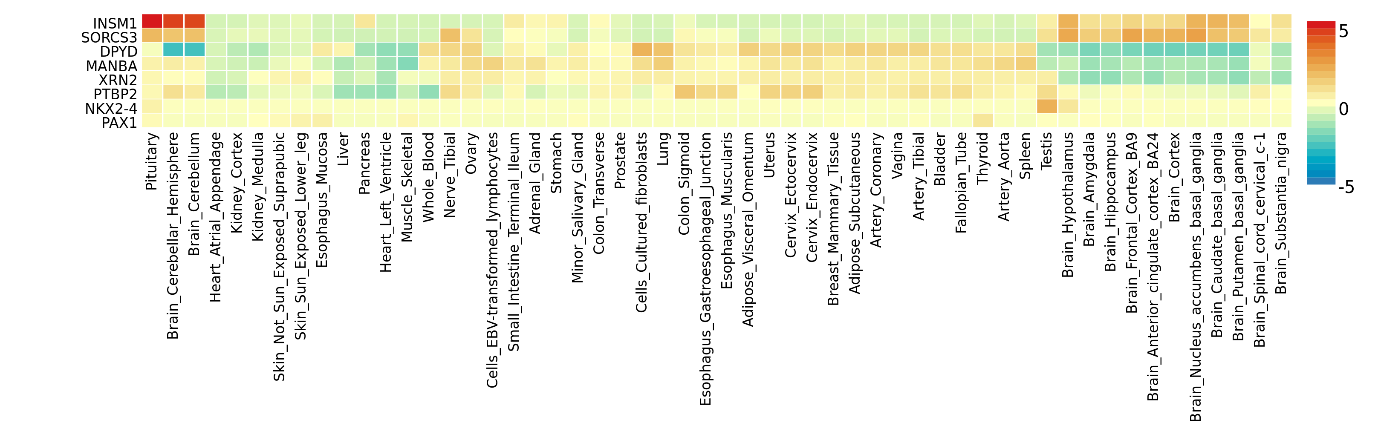

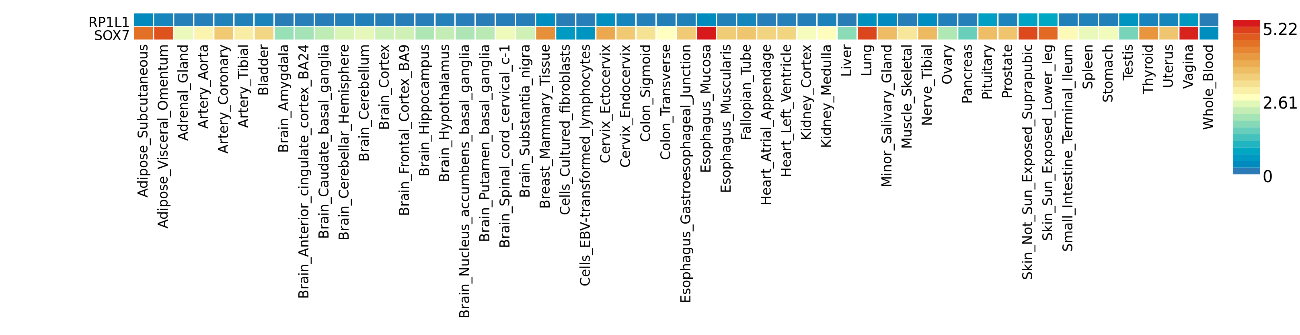

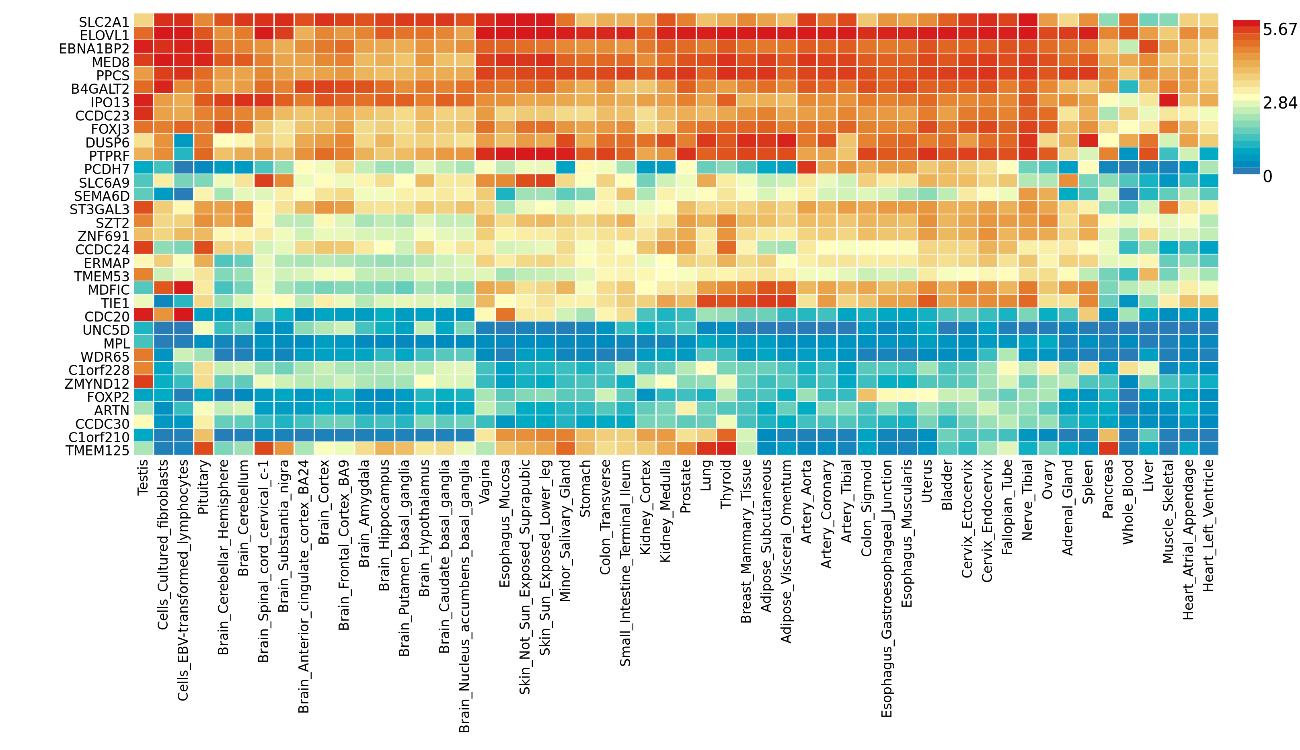

**Supp. Table 1.** FUMA analysis of SNPs shared by ASD and ADHD.

| ensg | symbol | Reported by Demontis et al. | Reported by Grove et al. or Alonso-Gonzalez et al.^#^ | CHR | Positional  mapping | EQTL  mapping | Chromatin  interaction  mapping | IndSigSNPs |
| --- | --- | --- | --- | --- | --- | --- | --- | --- |
| ENSG00000117569 | PTBP2 |  | X | 1 |  |  | X | rs2391769 |
| ENSG00000188641 | DPYD |  |  | 1 |  |  | X | rs2391769 |
| ENSG00000109323 | MANBA |  |  | 4 | X | X |  | rs227378 |
| ENSG00000156395 | SORCS3 | X |  | 10 | X |  | X | rs6584649 |
| ENSG00000173404 | INSM1 |  |  | 20 |  |  | X | rs6047319 |
| ENSG00000088930 | XRN2 |  | X | 20 | X |  | X | rs6047319 |
| ENSG00000125816 | NKX2-4 |  | X^#^ | 20 |  |  | X | rs6047319 |
| ENSG00000125813 | PAX1 |  |  | 20 |  |  | X | rs6047319 |

**Supp. Table 2.** FUMA analysis of SNPs specific to ASD.

| ensg | symbol | Reported by Grove et al. | CHR | Positional  mapping | EQTL  mapping | Chromatin  interaction  mapping | IndSigSNPs |
| --- | --- | --- | --- | --- | --- | --- | --- |
| ENSG00000183638 | RP1L1 | X | 8 | X |  |  | rs10099100 |
| ENSG00000171056 | SOX7 | X | 8 | X |  |  | rs10099100 |

**Supp. Table 3.** FUMA analysis of SNPs specific to ADHD.

| ensg | symbol | Reported by Demontis at al. | CHR | Positional | EQTL | Chromatin | IndSigSNPs |
| --- | --- | --- | --- | --- | --- | --- | --- |
|  |  |  |  | mapping | mapping | interaction |  |
|  |  |  |  |  |  | mapping |  |
| ENSG00000066056 | ARTN | X | 1 | X | X | X | rs37452; rs10890255; rs72678674; rs34898652; rs113551349; rs72678674:rs10890255:rs37452:rs34898652 |
| ENSG00000066135 | ATP6V0B | X | 1 | X |  |  | rs37452 |
| ENSG00000066185 | B4GALT2 | X | 1 | X |  |  | rs37452 |
| ENSG00000066322 | C1orf210 |  | 1 | X |  | X | rs2275180; rs2275180:rs10890255:rs34898652:rs72678674:rs37452 |
| ENSG00000117394 | C1orf228 |  | 1 |  |  | X | rs2275180:rs72678674:rs10890255:rs37452 |
| ENSG00000117395 | CCDC23 |  | 1 |  |  | X | rs2275180 |
| ENSG00000117399 | CCDC24 | X | 1 | X |  | X | rs37452; rs72678674:rs10890255 |
| ENSG00000117400 | CCDC30 |  | 1 |  |  | X | rs2275180 |
| ENSG00000117407 | CDC20 |  | 1 | X |  | X | rs2275180 |
| ENSG00000117408 | DPH2 |  | 1 | X |  |  | rs37452 |
| ENSG00000117410 | DUSP6 | X | 12 | X |  |  | rs704067 |
| ENSG00000117411 | EBNA1BP2 | X | 1 |  |  | X | rs2275180 |
| ENSG00000126091 | ELOVL1 |  | 1 | X |  | X | rs2275180; rs2275180:rs10890255:rs34898652:rs72678674 |
| ENSG00000126106 | ERMAP |  | 1 |  |  | X | rs2275180 |
| ENSG00000127125 | FOXJ3 |  | 1 |  |  | X | rs2275180:rs72678674:rs10890255 |
| ENSG00000128573 | FOXP2 |  | 7 | X |  | X | rs10261780 |
| ENSG00000132768 | HYI | X | 1 | X |  |  | rs2275180 |
| ENSG00000135272 | IPO13 |  | 1 | X |  | X | rs37452; rs72678674:rs10890255:rs37452 |
| ENSG00000137872 | KDM4A | X | 1 | X |  |  | rs10890255; rs34898652; rs72678674 |
| ENSG00000139318 | MDFIC |  | 7 |  |  | X | rs10261780 |
| ENSG00000142949 | MED8 | X | 1 | X | X | X | rs2275180; rs2275180:rs10890255:rs34898652:rs72678674:rs37452 |
| ENSG00000156687 | MPL |  | 1 | X |  |  | rs2275180 |
| ENSG00000159214 | PCDH7 |  | 4 | X |  |  | rs28411770 |
| ENSG00000159479 | PPCS |  | 1 |  |  | X | rs2275180 |
| ENSG00000164010 | PTPRF | X | 1 | X |  | X | rs34898652; rs10890255; rs72678674; rs2275180:rs10890255:rs34898652:rs72678674:rs37452:rs72678674 |
| ENSG00000164011 | SEMA6D | X | 15 | X |  |  | rs8039398 |
| ENSG00000169851 | SLC2A1 |  | 1 |  |  | X | rs2275180:rs72678674:rs10890255 |
| ENSG00000177868 | SLC6A9 | X | 1 | X |  |  | rs37452; rs113551349 |
| ENSG00000178922 | ST3GAL3 | X | 1 | X | X | X | rs72678674; rs10890255; rs37452; rs113551349; rs2275180:rs10890255:rs34898652:rs37452:rs72678674; rs34898652 |
| ENSG00000179178 | SZT2 | X | 1 | X | X | X | rs2275180; rs10890255; rs2275180:rs10890255:rs34898652:rs72678674:rs37452 |
| ENSG00000186409 | TIE1 |  | 1 | X | X | X | rs2275180; rs2275180:rs72678674:rs10890255 |
| ENSG00000196517 | TMEM125 |  | 1 |  |  | X | rs2275180 |
| ENSG00000198198 | TMEM53 |  | 1 |  |  | X | rs2275180:rs72678674:rs10890255:rs37452 |
| ENSG00000198520 | UNC5D |  | 8 |  |  | X | rs118041269 |
| ENSG00000198815 | WDR65 | X | 1 |  |  | X | rs2275180 |
| ENSG00000243710 | ZMYND12 |  | 1 |  |  | X | rs2275180 |
| ENSG00000253313 | ZNF691 |  | 1 |  |  | X | rs2275180 |

**Supp. Table 4.** Bidirectional Mendelian randomization (MR). Causal relationship between ASD as exposure (*p*-value at 1e-6) and ADHD as outcome (left). Causal relationship between ADHD as exposure (*p*-value at 1e-6) and ASD as outcome (right).

|  | ASD on ADHD | | | | | | |  | ADHD on ASD | | | | | | |
| --- | --- | --- | --- | --- | --- | --- | --- | --- | --- | --- | --- | --- | --- | --- | --- |
|  | n(SNPs)=15 | | | | | | |  | n(SNPs)=41 | | | | | | |
|  | β |  | 95%-CI |  | SE |  | *p*-value |  | β |  | 95%-CI |  | SE |  | *p*-value |
| MR-IVW | 0.51 |  | (0.33; 0.69) |  | 0.09 |  | <0.001 |  | 0.34 |  | (0.24; 0.43) |  | 0.05 |  | <0.001 |
| MR weight median | 0.57 |  | (0.40; 0.74) |  | 0.09 |  | <0.001 |  | 0.36 |  | (0.26; 0.45) |  | 0.05 |  | <0.001 |
| MR weight mode | 0.56 |  | (0.17; 0.94) |  | 0.20 |  | 0.013 |  | 0.35 |  | (0.14; 0.57) |  | 0.11 |  | 0.003 |
| MR-RAPS | 0.56 |  | (0.40; 0.72) |  | 0.08 |  | <0.001 |  | 0.35 |  | (0.24; 0.45) |  | 0.05 |  | <0.001 |
| MR-Egger | 0.76 |  | (0.10; 1.42) |  | 0.34 |  | 0.025 |  | 0.06 |  | (-0.35; 0.48) |  | 0.21 |  | 0.8 |
| *MR-Egger intercept* | *-0.02* |  | *(-0.08; 0.03)* |  | *0.03* |  | *0.5* |  | *0.02* |  | *(-0.01; 0.06)* |  | *0.02* |  | *0.19* |
| MR: Mendelian randomization. SNPs: Single-Nucleotide Polymorphisms. ASD: Autism Spectrum Disorder. ADHD: Attention Deficit Hyperactivity Disorder. | | | | | | | | | | | | | | | |
